# Phyllosphere and rhizosphere microbiomes empower *Nicotiana tobacum* complex traits dissection and prediction

**DOI:** 10.64898/2026.02.22.707303

**Authors:** Qiao Du, Yu Han, Zhengwen Liu, Huan Si, Yan Ji, Li Liu, Zhiliang Xiao, Lirui Cheng, Aiguo Yang, Dan Liu, Yanjun Zan

## Abstract

Understanding how plant-associated microbiomes interact with host genome variation to influence agronomic traits is essential for advancing microbiome⍰assisted crop improvement. In this study, we characterized the phyllosphere and rhizosphere microbiomes of 164 diverse *Nicotiana tabacum* accessions using 16S rRNA sequencing and integrated these data with host genomic variation and 22 agronomic traits. The two microbiomes exhibited distinct taxonomic structures, diversity patterns, and predicted metabolic functions. Microbiome genome⍰wide association studies identified extensive host genetic control over microbial abundance, including 49 shared genomic loci that explained nearly half of the heritable variation in both microbiomes. Microbiome⍰wide association studies revealed biologically meaningful associations between specific ASVs and agronomic traits. However, network analysis demonstrated that microbial sub⍰communities, rather than individual taxa, contributed substantially to phenotypic variation. Then, colocalization analysis further identified genetic variants jointly influencing microbial abundance and metabolite traits, highlighting potential host-microbe-trait causal links. Incorporating microbiome data into genomic selection models, we successfully improved prediction accuracy for several traits, especially plant architecture and flowering. Together, this work provides a comprehensive population⍰level framework linking host genetics, microbiome composition, and agronomic traits in tobacco, offering new insights for microbiome⍰informed breeding strategies.

## Introduction

Plants harbor diverse microorganisms across various tissues, forming highly structured taxonomic communities(3). These microorganisms establish symbiotic relationships with plants and play pivotal roles in enhancing plant fitness in natural environments, specifically by improving growth, stress tolerance, pathogen resistance, and nutrient acquisition (4–8). Consequently, modulating the plant-associated microbiome represents a promising frontier for enhancing agricultural yields and mitigating diseases (9,10). However, microbial communities are complex, shaped by the interplay between host, environmental conditions, and inter-microbe interactions (11). While recent studies have shown that microbiome composition is significantly influenced by host genetic effects (12–15), the intricate relationship among host genetic variations, the plant-associated microbiome, and agronomic traits variation remains poorly characterized.

Genome-wide association study (GWAS) provide a powerful, hypothesis-free approach to elucidate the link between genetic variation and complex traits variation (16) in important crops, such as rice, potato, and tomato (17–20). Recently, this framework has been extended to treat the entire microbial community as a complex, heritable trait. This microbiome-based GWAS (mGWAS) approach enables the detection of associations between host genetic loci and microbial composition (12,21–25). For instance, GWAS of the *Arabidopsis thaliana* phyllosphere microbes identified host loci involved in defense and cell wall integrity (22). Similarly, a study on the rhizoplane microbiota of foxtail millet demonstrated that microbial composition is affected by allelic variations in genes related to immunity, metabolites, hormone signaling, and nutrient uptake (21). While these studies validate the utility of GWAS for host-microbiome genetics, the genetic basis of host-microorganism interactions remains largely unexplored. Specifically, how these interactions translate into host phenotypic traits variation is not well understood, limiting the utilization of microbial engineering for targeted agronomic improvement. Furthermore, because the impacts of microorganisms tend to be specific to plant species and cultivars (26,27), investigating these dynamics in diverse genetic backgrounds is essential.

*Nicotiana tabacum* (common tobacco) is an important economic crop. Beyond its commercial value, N. tabacum is widely utilized as a model plant for genetic and biological studies due to its high transformation efficiency and extensive phenotypic diversity (29–31). Previous research utilizing quantitative trait loci (QTLs) has identified multiple loci linked to key agronomic traits, such as leaf morphology, nicotine content, plant height, and pathogen defense (32–34). The recent publication of a chromosome-level genome assembly for *N. tabacum* has further empowered the dissection of complex trait evolution and genetic regulation (35).

In this study, we profiled the rhizosphere and phyllosphere microbiomes of 164 diverse N. tabacum accessions using 16S rRNA sequencing. We performed microbiome genome-wide association studies (mGWAS) to uncover the host genetic variations that shape microbial composition. Leveraging a comprehensive dataset of 22 agronomic traits, we subsequently identified genetic loci and specific microbes influencing host phenotype through integrated GWAS and microbiome-wide association studies (MWAS). Moreover, we demonstrated the efficacy of incorporating microbiome data into genomic selection models to enhance trait prediction. This work establishes the reciprocal interactions among host genetic variation, tobacco-associated microbiota, and agronomic traits, offering promising insights into new strategies for improving tobacco productivity through microbiome engineering.

## Materials and Methods

### Plant cultivation and sampling

A total of 164 germplasm resources that are preserved in the GeneBank of Tobacco hosted at the Tobacco Research Institute, Chinese Academy of Agriculture Sciences were selected in our study.

At the year of 2023, we started to phenotype the collection at Qingdao, China (120.45° E, 36.38° N) with a randomized complete block design with two replicates. One replication of each line includes two rows, each with 10 replicates of the same genotype. Each row is 10 meters in length with a row spacing of 1.2 m and a plant distance of 0.5 m. Detailed description on traits measurements were mentioned in previous studies12,13,87. Briefly speaking, flowering time related traits were obtained by individually counting the days to budding and flowering. Plant architecture traits were manually measured for each plant using ruler and angle ruler. Two mature middle leaves from each plant were harvested and measured for leaf morphological traits and metabolic traits. All phenotypes were measured right before the first flower blooms and averaged within each replicated row. Averaged measurements among ten replicates were adjusted using a linear model, fitting blocks and rows as fixed effects, to calibrate the spatial variation, generating the phenotype records for downstream analysis. Three replicates of roots and leaf were sampled during flowering for 16s Sequencing at Novogene.

### DNA extraction, PCR amplification, and 16s rRNA sequencing

Genomic DNA was extracted from all samples and quantified using a Nanodrop spectrophotometer. Amplification of the V3–V4 region of the 16S rRNA gene was performed. The amplicons were purified using magnetic beads. Purified products were quantified by fluorescence-based methods and then pooled. The library was prepared using the VAHTS Universal DNA Library Prep Kit for Illumina V3 and VAHTS DNA Adapters set3–set6 for Illumina (Vazyme Biotech Co., Ltd, Nanjing, China). Quality assessment was conducted on the constructed library, and qualified libraries were subjected to high-throughput sequencing on the Illumina NovaSeq 6000 platform.

### 16s rRNA sequencing analysis

Raw sequences were filtered and demultiplexed based on unique barcode sequences, followed by removal of barcodes and primer sequences. Following these preprocessing steps, we obtained an average of 121,035 and 125,072 clean reads from the rhizosphere and phyllosphere per sample. Sequences denoising was performed using the DADA2 plugin (36) within the QIIME2 pipeline (37) to generate high-quality amplicon sequence variants (ASVs). Taxonomy was assigned to ASVs using the QIIME2 classify-sklearn plugin (38) with a Naïve Bayes classifier. The SILVA database (Release 138) (39) was used for classifier training and classification.

For downstream analyses, we focus on 765 high-abundance ASVs, defined as those with at least 3 reads in at least 50% of the samples. To account for varying sequencing depths, samples were rarefied at a threshold of minimum sequence frequency across all samples. The abundances of ASVs were then normalized to relative abundance. Alpha-diversity analysis with Shannon and Simpson indexes was performed by R package ‘vegan’ (40) with Wilcoxon’s rank-sum test. Beta diversity, based on Bray-Curtis distances, was visualized using principal coordinates analysis (PCoA), and the statistical significance of group separation was assessed by PERMANOVA with the adonis function (R package ‘vegan’). The stacked column chart and Linear Discriminant Analysis Effect Size (LEfSe) were completed using Wekemo BioinCloud. The LEfSe analysis (41) utilized the Kruskal–Wallis test and LDA score to identify differentially abundant taxa between groups.

Functional potential was inferred from 16S rRNA data using PICRUSt2 via the Wekemo BioinCloud platform. Differentially abundant metabolic pathways were identified using STAMP (v2.1.3) based on Welch’s t-test, with $p$-values adjusted for multiple testing using the Benjamini–Hochberg False Discovery Rate (FDR < 0.05). Finally, Mantel tests were employed to evaluate the correlation between host genetic distances and microbiome distances using the ‘vegan’ package.

### Whole-genome sequencing and SNP calling

All samples were sequenced on the Illumina HiSeq X Ten platform with 150 bp paired end reads. After quality control, the clean reads were aligned to the *N.tabacum* reference genome using BWA-MEM (35,42). The aligned reads were processed with SAMtools to sort and remove duplicated reads (43). SNP calling was performed using freebayes with default parameters (44). For each sample, SNPs were filtered with the following parameters: QUAL>20, DP>3, F_MISSING<0.5, and -m2 -M2 by VCFtools (45). Then, SNPs were merged across all samples and further filtered with a missing rate < 0.5. Beagle was used to impute missing genotypes (46). Finally, MAF > 0.05 and pairwise r^2^< 0.95 in 10 Mb sliding windows were applied to obtain 1,465,715 high-quality SNPs for downstream analysis.

### GWAS analysis on microbiome and agronomic traits

Before GWAS analysis, the SNP-based kinship heritability of each ASV and agronomic trait was estimated using the GREML-AI method in GCTA software (47). GWAS were then performed in GCTA using the mixed linear model (MLM) to test the associations between SNPs and microbial ASVs abundance and agronomic traits. The significant threshold was set as Bonferroni correction (p < 0.05/number of SNPs). For partitioning phenotypic variance explained (PVE), SNPs were split into two sets: the 49 shared-locus SNPs and the remaining genome-wide SNPs. We computed separate genetic relationship matrices (GRMs) for each SNP set and used GREML in GCTA to estimate the PVE attributable to each GRM by quantifying the relative contribution of the shared loci by the ratio of the genetic variance from the shared-locus GRM to the sum of genetic variances from both GRMs.

### MWAS analysis on agronomic traits

To evaluate the association between microbial ASVs and agronomic traits, we performed microbiome-wide association studies (MWAS) using a general linear model (GLM) implemented in the R package ‘lm’. The top 10 principal components (PCs) calculated from the SNP data were included as covariates to control for population structure. The significant threshold was corrected by the Bonferroni method (p < 0.05/number of ASVs).

### Microbial co-occurrence network construction and contribution analysis

A microbial co-occurrence network was constructed by the R package ‘microeco’ (48). The ASVs data table of rhizosphere and phyllosphere was merged, and Spearman rank correlation coefficients were calculated for all ASV pairs. Bonferroni-corrected p-values were used to identify significant correlations and construct the connection edges between ASVs. The resulting network was partitioned into 29 sub-communities using the fast-greedy modularity optimization algorithm.

After identifying the sub-communities, we assessed their contribution to agronomic traits. For each trait, we calculated the PVE by ASVs within each sub-community using a generalized linear model with the top 10 genetic PCs as covariates. PVE was calculated as the ratio of the variance explained by the ASVs in the sub-community to the total phenotypic variance. To compare the PVE of sub-communities with individual ASVs, we also calculated the PVE for each ASV on the same trait using the same model. To control for the potential bias of increased predictor variables and confirm that the observed PVE was driven by biologically meaningful sub-community structures, we performed a permutation analysis. Specifically, for each sub⍰community, we randomly shuffled the ASV abundance values across samples while keeping the same set of ASVs and the same model structure 100 times. We then recalculated the PVE using the permuted data. This procedure preserves the number and identity of ASVs but disrupts their ecological coherence.

### Colocalization analysis

To identify genetic loci simultaneously associated with microbial ASVs and agronomic traits, we performed colocalization analysis using the R package ‘coloc’ (49). For each pair of microbial ASV and agronomic trait that shared a significant GWAS locus, we extracted summary statistics (effect sizes, standard errors, allele frequencies) for all SNPs within a 1 Mb window centered on the lead SNP. The coloc.abf function was then used to calculate the posterior probabilities for five hypotheses: no association with either trait (H0), association with only the first trait (H1), association with only the second trait (H2), association with both traits but different causal variants (H3), and association with both traits and a shared causal variant (H4). A high posterior probability for H4 (PP4 > 0.9) was considered strong evidence for colocalization.

### Microbiome-integrated genomic selection analysis

The genomic selection analysis was performed using the mixed linear model by incorporating both genetic relationship matrix and microbial relationship matrix by fitting the models below:

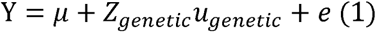

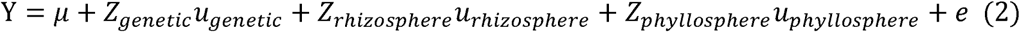

Where Y is the vector of phenotypic values, *µ* is the overall mean intercept, *z_genetic_*, *z_rhizosphere_* and *z_phyllosphere_* are incidence matrices relating observations to genetic effects *u_genetic_*, rhizosphere microbial effects *u_rhizosphere_*, and phyllosphere microbial effects *u_phyllosphere_*, respectively. The random effects are assumed to follow normal distributions:

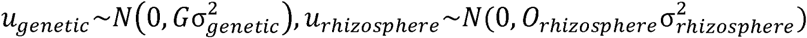

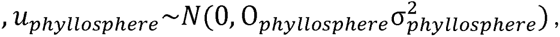 and 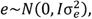 where G is the genetic relationship matrix calculated from SNP data by GCTA (107), *O_rhizosphere_* and *O_phyllosphere_* are the microbial relationship matrices calculated from rhizosphere and phyllosphere ASV abundance data by OSCA (50), respectively, I is the identity matrix, and 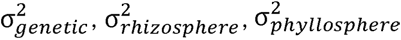 and 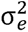 are the variances of genetic effects, rhizosphere microbial effects, phyllosphere microbial effects, and residuals, respectively.

For SNP only model (1), only genetic effects were considered, while for SNP + microbiome model (2), both genetic and microbial effects were included. Three-fold cross-validation was performed to evaluate the prediction accuracy of both genomic selection models. The dataset was randomly divided into three equal parts. In each iteration, two parts were used as the training set to fit the model and estimate the effects, while the remaining part served as the test set to predict phenotypic values. This process was repeated 100 times to ensure robustness, and the Pearson correlation coefficient between observed and predicted phenotypic values was calculated to assess prediction accuracy. The t-test was used to compare the prediction accuracies between the two models across all traits.

## Results

### Comparison of rhizosphere and phyllosphere microbial community composition for Nicotiana tabacum

In this study, 16S rRNA gene sequencing was performed for the rhizosphere and phyllosphere for 164 *Nicotiana tabacum* samples. The flat Rarefaction Curve indicated that the sequencing depth was sufficient to capture the full extent of bacterial diversity (Fig. S1). After quality control and taxonomic assignment (Materials and Methods), a total of 765 ASVs were annotated with 22 phyla, 89 orders, and 114 families.

Assessments of $\alpha$-diversity revealed significant differences between the two niches, with the rhizosphere microbiome exhibiting markedly higher Shannon and Simpson indices compared to the phyllosphere (*p* < 0.05; Fig. 1A). These results underscore the greater microbial richness and heterogeneity present within the rhizosphere environment. Principal Coordinates Analysis (PCoA) on Bray-Curtis distances at the orders-level demonstrated clear niche-based clustering, with rhizosphere and phyllosphere samples forming distinct groups (Fig. 1B). This separation confirms that the two microbial communities possess fundamentally different taxonomic structures and compositional profiles.

**Fig. 1.**
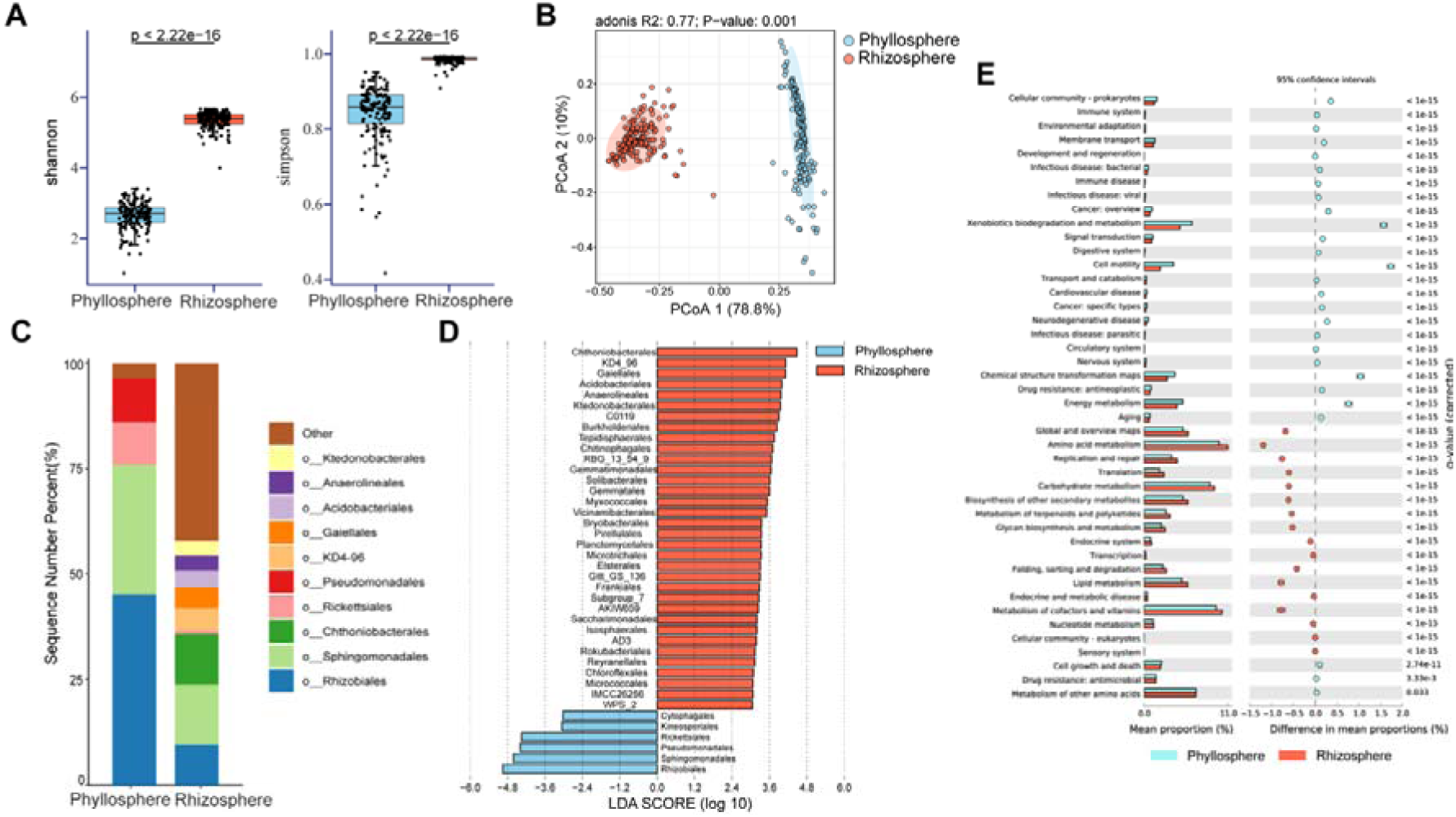
Analyses of microbial composition and function between the rhizosphere and phyllosphere. **(A)** Alpha diversity estimates (Shannon and Simpson indices) between two groups (Wilcoxon’s rank-sum test, *p* < 0.05). **(B)** PCoA plot based on Bray-Curtis distance of order-level relative abundance of microbiota between two groups (Adonis, R^2^ = 0.77, *p* = 0.001). **(C)** Top 10 abundant microbial orders in two groups. **(D)** LEfSe analysis of microbiota in two groups at the orders level (Kruskal–Wallis test; false discovery rate (FDR) <□0.05, LDA (Linear Discriminant Analysis) □>□3). **(E)** Functionally predicted pathways differ in abundance in two groups. The bar plot shows mean proportions of pathways. Only *p* < 0.05 is shown (Welch’s t-test, FDR-adjusted).

At the order level, the phyllosphere microbiome was dominated by Rhizobiales, Sphingomonadales, Pseudomonadales, and Rickettsiales (Fig. 1C). In contrast, the rhizosphere was primarily composed of Sphingomonadales, Chthoniobacterales, Rhizobiales, Gaiellales, and the class-level group KD4-96. In total, 40 taxonomic orders differed significantly between phyllosphere and rhizosphere (*p* < 0.05, Fig. 1D, Kruskal-Wallis tests). Specifically, six orders were significantly enriched in the phyllosphere including Pseudomonadales, Sphingomonadales, and Cytophagales, which are frequently associated with nutrient acquisition and polysaccharide metabolism (Fig. 1D). Conversely, the orders enriched in the rhizosphere—such as Acidobacteriales, Gaiellales, KD4-96, and Burkholderiales—are typically linked to soil-specific ecological functions.

Functional prediction revealed that 44 metabolic pathways differed significantly between the two microbial communities (*p* < 0.05, Fig. 1E). Of these, 17 pathways were significantly enriched in the rhizosphere, focusing on nutrient cycling and primary metabolism, such as the metabolism of amino acids, carbohydrates, nucleotides, and lipids. Conversely, the 27 pathways enriched in the phyllosphere were predominantly associated with host-microbe interactions and environmental adaptation, including stress response, xenobiotic biodegradation, signal transduction, and responses to infectious diseases.

### Genetic determination of the phyllosphere and rhizosphere microbiota composition

Most ASVs exhibited moderate kinship heritability, with median value of 0.28 and 0.15 for the phyllosphere and rhizosphere microbiomes, respectively (Fig. S2), , indicating a substantial host genetic influence on microbial recruitment. To elucidate the genetic basis of microbiome composition, we performed microbiome genome-wide association studies (mGWAS) on all phyllosphere and rhizosphere ASVs using a mixed linear model (Table S3, S4). Due to the extensive linkage disequilibrium (Fig. S3), significant SNPs were grouped into 1 Mb genomic bins to facilitate comparison and visualization (Materials and Methods).

A total of 80 phyllosphere and 516 rhizosphere ASVs were significantly associated with at least one SNP (Fig. 2A, B). Across 4,061 genomic bins, we identified 776 and 1,916 loci associated with phyllosphere and rhizosphere variation, respectively. Notably, 441 loci were shared between both niches (Fig. 2C), suggesting that pleiotropic genetic factors regulate microbial abundance across different plant tissues. Among these, 49 loci were consistently identified at least twice in both microbiomes. Furthermore, the number of associated ASVs in the two groups showed a strong positive correlation (Fig. 2D).

**Fig. 2.**
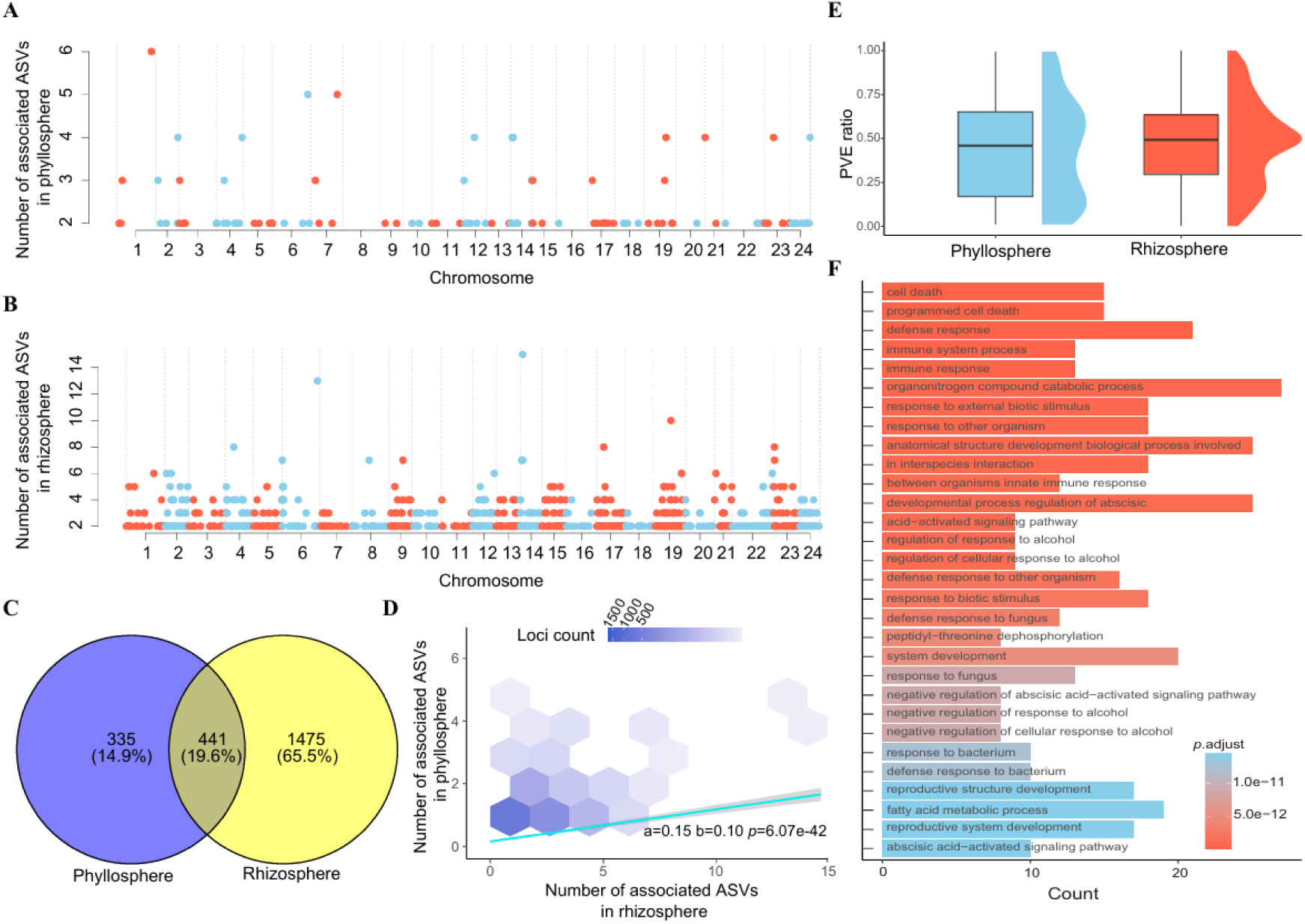
Genetic determination of the phyllosphere and rhizosphere microbial community composition. (A) Manhattan-like plot of the phyllosphere microbiome. Each point represents a genomic bin, with the height of each point indicating the number of ASVs significantly associated with SNPs within that bin. (B) Manhattan-like plot of rhizosphere microbiome. (C) Venn diagram showing the overlap of significantly associated genomic bins between the phyllosphere and rhizosphere microbiomes. (D) Hexbin scatter plot comparing the number of associated ASVs in each bin between the phyllosphere and rhizosphere microbiomes. Each hexbin represents a data area, and its color displays the number of loci in this area. The regression line shows an intercept of 0.15, a slope of 0.10, and a *p*-value of 6.07×10^-42^. (E) Box plot illustrating the proportion of ASV variation in the phyllosphere and rhizosphere microbiomes explained by SNPs from the 49 common genomic bins. (F) The most significant 30 GO terms related to the biological processes of the genetic loci located at chromosome 14: 12,992,799–13,992,799 bp.

Importantly, the SNPs within these 49 shared loci—representing only 1.2% of the genome and 2.0% of the total SNP set—accounted for approximately half of the total heritable variation (median PVE: phyllosphere = 0.46; rhizosphere = 0.49; Fig. 2E). This highlights the presence of genetic “hotspots” that play a vital role in regulating microbial abundance. For instance, a locus on chromosome 14 (12.99–13.99 Mb) exhibited the highest number of associations, linking to 4 phyllosphere and 15 rhizosphere ASVs. This locus contains 21 genes enriched in Gene Ontology (GO) terms related to plant immunity and microbial interactions, including responses to external biotic stimuli, fungi, and bacteria (Fig. 2F).

### Microbiota composition is associated with agronomic traits variation

To investigate the influence of the microbiome on host performance, we performed association analyses between microbial composition and 22 agronomic traits. These traits, which exhibited a median heritability (*h^2^*) of 0.66 (Table S5), were categorized into five groups: plant architecture, leaf architecture, disease resistance, flowering time, and metabolites (Fig. S4). Mantel tests revealed significant correlations between microbial community structure and host phenotypes. Notably, the phyllosphere microbiome showed a stronger association with plant architecture, whereas the rhizosphere microbiome was primarily linked to metabolites and disease resistance (Fig. 3A).

**Fig. 3.**
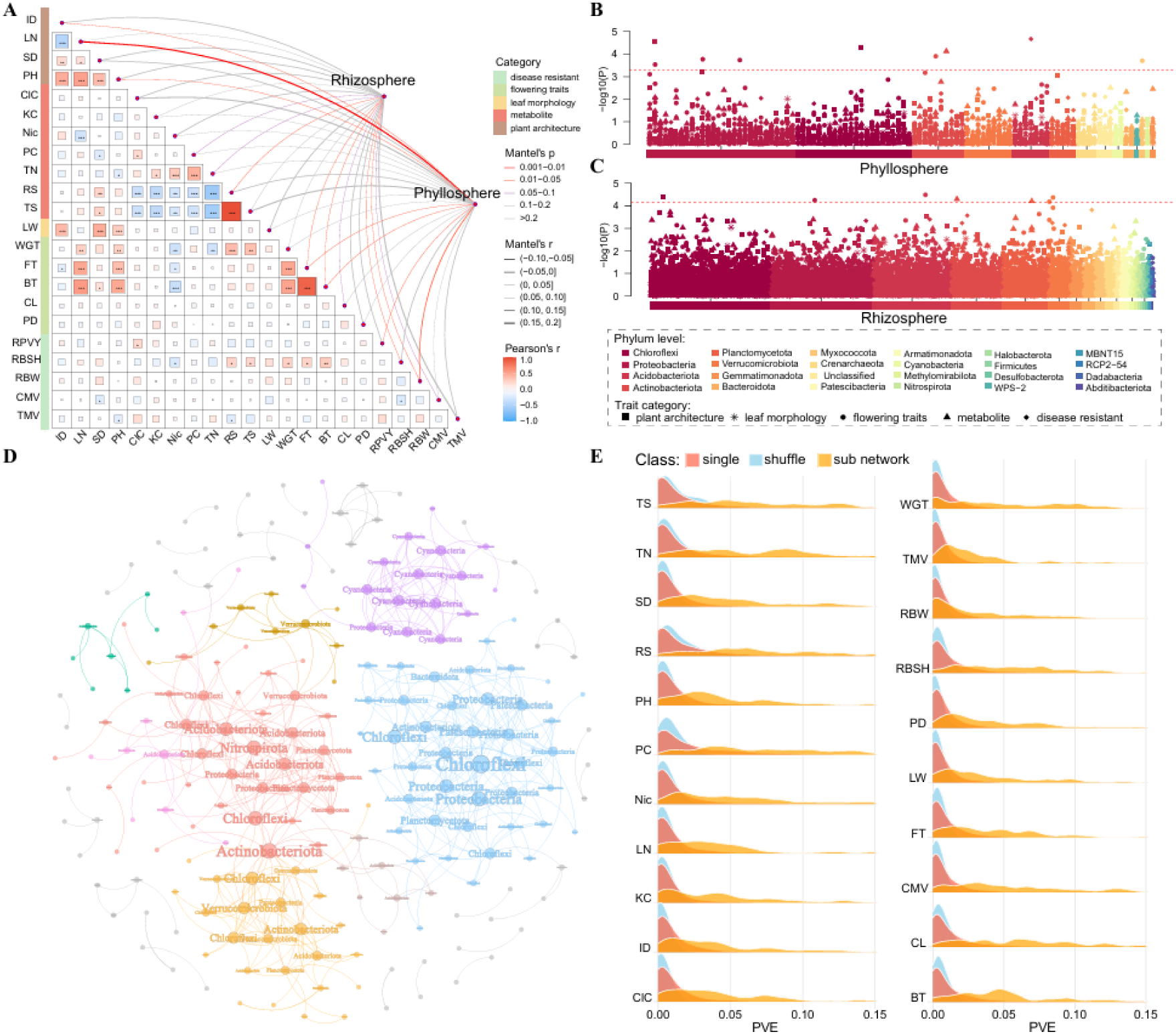
Microbial community composition is associated with agronomic traits variation. **(A)** Mantel test revealed association between different microbiomes and agronomic traits. The lower triangle heatmap represents Pairwise Pearson correlation coefficients for agronomic traits. The width of the edges corresponds to the Mantel correlation coefficient, and the color represents the significance. Manhattan-like plots of MWAS (Microbiome-Wide Association Study) in the phyllosphere (**B**) and rhizosphere (**C**) microbiomes. Each point represents an ASV, with the color indicating the phylum and the shape representing the associated trait category. **(D)** Network analysis of the microbial community. Different colors of edges and dots represent distinct sub-communities. The size of the dots corresponds to the number of connections for each ASV. **(E)** Density plots of the proportion of variance explained (PVE) by ASVs across all traits. The height of each plot reflects the statistical distribution of each class, with higher values indicating greater frequency. The red density plots display the PVE distribution of individual ASVs, the orange density plots show the PVE distribution of ASVs within sub-communities, and the blue density plots represent the PVE distribution of shuffled-abundance ASVs within sub-communities.

To identify specific ASVs linked to these phenotypes, we conducted microbiome-wide association studies (MWAS). Under a rigorous Bonferroni-corrected threshold, only 9 phyllosphere and 7 rhizosphere ASVs were significantly associated with agronomic traits (Fig. 3B, C; Table S6). Among these, flowering traits exhibited the highest degree of association, linking to nine specific ASVs. A noteworthy finding was ASV5 (Methylobacterium), which was associated with both vegetative growth (plant height) and reproductive onset (budding time). This association is mechanistically plausible, as Methylobacterium species are known to produce phytohormones—such as cytokinins and indole-3-acetic acid—that promote biomass and height in diverse plant species (51,52).

However, the limited number of significant associations (0.09% of all combinations) suggests that the independent impact of individual microbes on complex trait variation is marginal. This is consistent with the highly polygenic nature of these traits. Consequently, we shifted our focus from individual ASVs to the collective effect of microbial sub-communities. A co-occurrence network was constructed (Fig. 3D; Materials and Methods), identifying 211 ASVs with 509 significant connections. While the network revealed high connectivity among ASVs from different phyla, no connections were observed between phyllosphere and rhizosphere communities (Fig. S5), reaffirming their distinct ecological structures. The network can be partitioned into 29 sub-communities, with 155 ASVs forming eight primary modules (Fig. 3D; Table S7). Crucially, these sub-communities explained a significantly higher proportion of phenotypic variance (PVE) across all traits compared to individual ASVs or randomly shuffled null models (Fig. 3E). Collectively, these results underscore that agronomic traits are primarily shaped by collective microbial action rather than the influence of isolated taxa.

### Colocalization of genetic variants associated with microbial and agronomic traits

To detect genetic variants simultaneously associated with agronomic traits and microbiome composition, we performed GWAS on all 22 agronomic traits. This analysis identified 59 loci significantly associated with agronomic traits (Fig. 4A). Notably, a majority (54%) of these loci were also identified in our rhizosphere mGWAS, while 24% overlapped with loci from the phyllosphere mGWAS (Fig. 4B). The high degree of overlap suggests a shared genetic architecture governing both host traits and their associated microbial communities. Among these shared loci, colocalization analysis (Fig. 4C) identified a high-confidence locus associated with ASV141 (KD4-96, PP4 = 0.96) in the rhizosphere and leaf total reducing sugar level (Fig. 4D, E). This finding suggests that the identified genetic variant exerts a pleiotropic effect, simultaneously regulating tobacco sugar content and shaping the abundance of KD4-96 Members of the KD4-96 group are recognized for their roles in herbicide biodegradation, heavy metal tolerance, and nutrient cycling (53–55). By modulating rhizosphere nutrient-toxic element dynamics and influencing plant metal uptake, KD4-96 has been shown to promote plant growth and enhance crop yield (56,57).

**Fig. 4.**
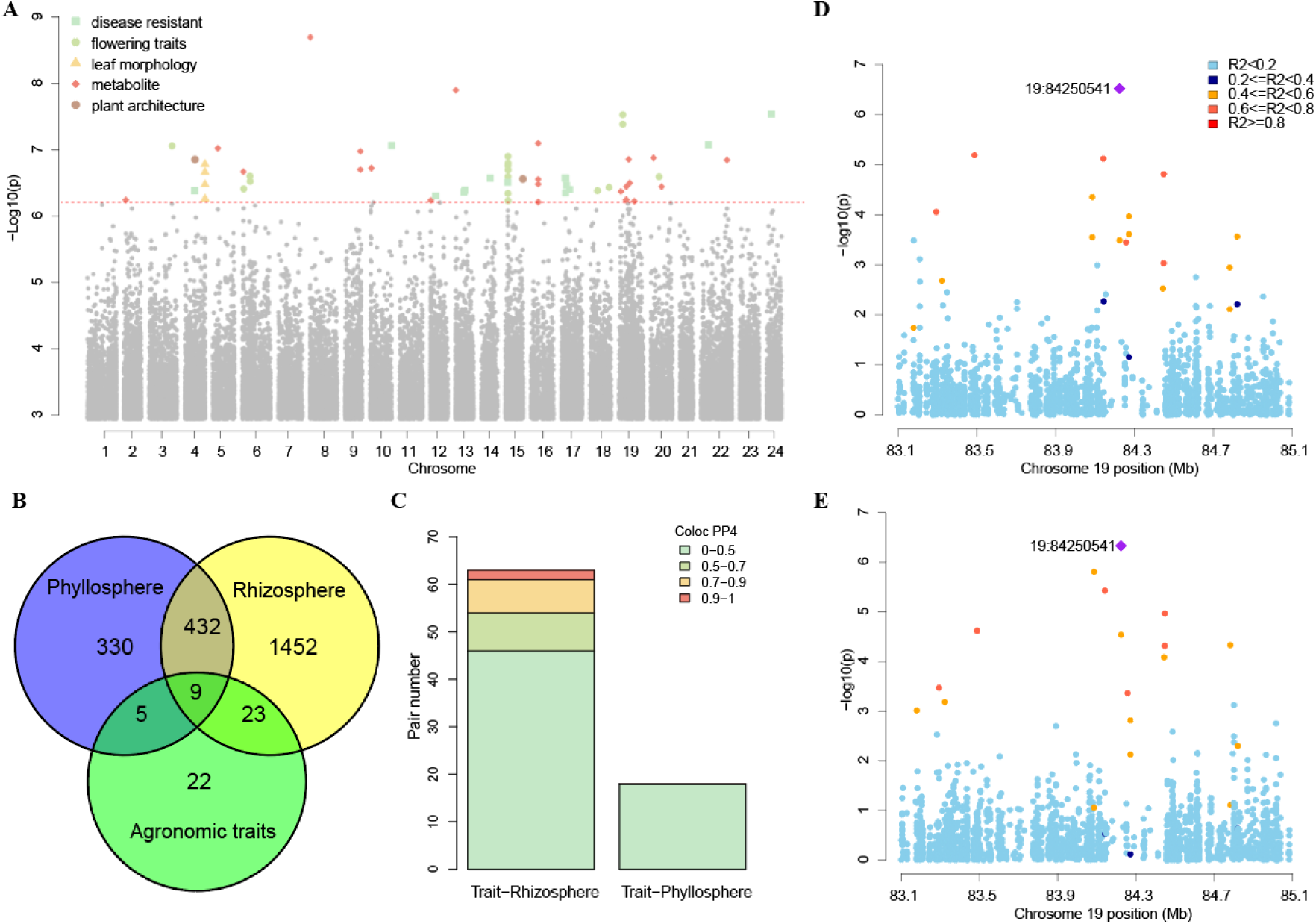
Genetic variants associated with agronomic traits and microbiome composition. **(A)** Joint Manhattan plots overlaying results from all agronomic traits. The red horizontal dashed lines are the Bonferroni corrected threshold. Significant SNPs were marked with corresponding trait types. **(B)** Venn diagram showing the overlap of significantly associated genomic bins between the phyllosphere and rhizosphere microbiomes and agronomic traits. **(C)** The Posterior Probability 4 (PP4) of each colocalization analyzed pairs. PP4 represents the possibility that signals observed in two different genetic association studies arise from the same genetic variants. The bars are colored by the different stringency levels for colocalization PP4. Example of colocalization (PP4=0.96) for a mGWAS locus associated with ASV141 in the rhizosphere **(D)** and a GWAS locus associated with total reducing sugar **(E)** on chromosome 19. The top colocalized SNP (19: 84250541) was marked as a purple triangle.

**Fig 5.**
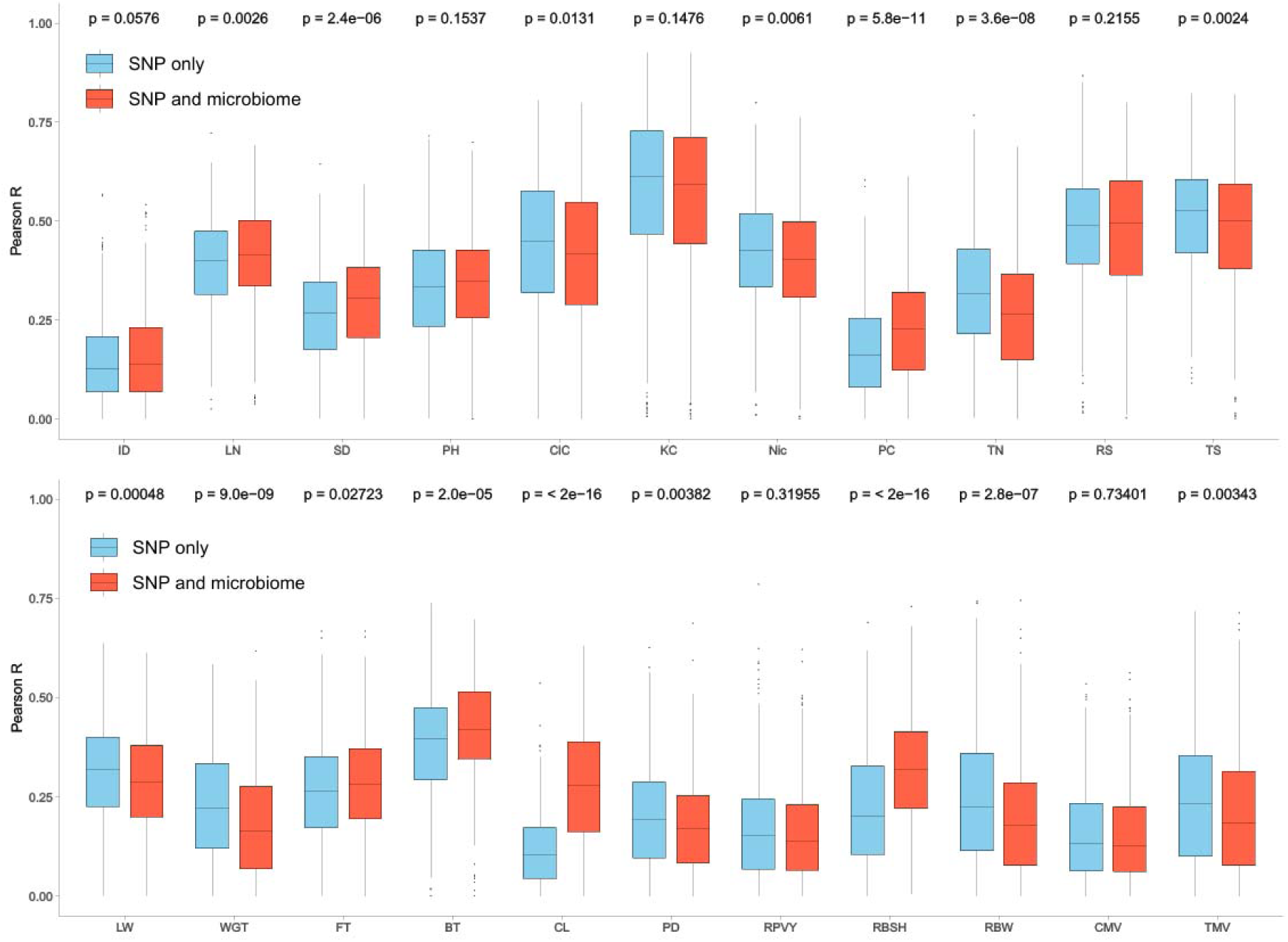
Genome selection accuracy results. The boxplot illustrates the 5-fold cross-validation results of the Pearson correlation between raw phenotype and predicted phenotype. Each validation was repeated 100 times to ensure an even sample of train and test individuals.

### Microbiomes improve genome selection accuracy

Genome selection (GS) is a promising technique in agronomic breeding, , enhancing efficiency by predicting phenotypes based on whole-genome variation rather than relying solely on labor-intensive phenotypic evaluations. While traditional GS models utilize only genotype data, the integration of multi-layered data—such as environmental factors and various “omics” datasets—can significantly enhance prediction accuracy (58). Using Genomic Best Linear Unbiased Prediction (GBLUP), we assessed whether incorporating microbiome data could improve the prediction of 22 tobacco agronomic traits.

Compared to models using genotype data alone, the inclusion of microbiome data significantly improved prediction accuracy for eight traits, showed no significant change for five traits, and resulted in a slight reduction for nine traits. Notably, traits related to plant architecture and flowering time exhibited the strongest potential for improvement. These results suggest that integrating microbiome data into breeding programs is particularly effective for predicting complex morphological and phenological traits, offering a novel strategy for microbiome-enhanced crop improvement.

## Discussion

In this study, we characterized the rhizosphere and phyllosphere microbiomes of tobacco, identifying significant differences in community diversity, taxonomic composition, and predicted metabolic functions. Our findings reinforce the established paradigm that distinct plant compartments serve as unique ecological niches, selectively recruiting specialized microbiota (59–62). Specifically, the significantly higher α-diversity observed in the rhizosphere compared to the phyllosphere aligns with the broader understanding that belowground environments—influenced by complex soil matrices and root exudates—harbor richer microbial communities than the more exposed and nutrient-limited aboveground niches (60,62,63).

The composition of these microbiota was markedly distinct between the two nitches. In the phyllosphere, we found a significant enrichment of Rhizobiales, Sphingomonadales, Pseudomonadales, and Rickettsiales, which are consistent with dominant taxa reported in other major crops (64). For instance, in wheat, these groups are linked to organic acid metabolism and nitrogen input, suggesting a conserved role in mediating plant nutrient responses (65). Interestingly, Rhizobiales and Sphingomonadales also maintained dominance within the tobacco rhizosphere. Members of the order Rhizobiales are well-documented plant growth-promoting bacteria that facilitate nutrient uptake (66), while many Sphingomonadales species enhance growth and stress tolerance (67,68). The prevalence of these core taxa across both niches suggests they fulfill fundamental ecological roles essential for tobacco fitness, regardless of the tissue-specific environment.

In the rhizosphere, the enrichment of Gaiellales and the KD4-96 group further characterizes the niche’s functional profile. Gaiellales are known to decompose soil organic matter and facilitate phosphate acquisition (69),while KD4-96 has been positively correlated with enhanced plant growth and yield (56,57). Additionally, the enrichment of Acidobacteriales and Burkholderiales suggests a community geared toward biogeochemical cycling and plant health promotion (70) (71). Our functional inference analysis supports this niche differentiation: while the rhizosphere focuses on the degradation and conversion of organic substrates, the phyllosphere is specialized for environmental stress adaptation and signal transduction. This aligns with the understanding that leaf microbiota are frontline players in plant defense and atmospheric adaptation (73,74)

While host genetics have been shown to shape the microbiome in crops like maize and rice (21,23,24,75), our study provides the first high-resolution map of the tobacco microbiome’s genetic architecture. Despite the distinct ecological pressures of the rhizosphere and phyllosphere, the discovery of partly shared genetic loci highlights a concentrated host mechanism that regulates microbial communities across different tissues. For instance, the shared locus on chromosome 14, enriched with immune-related and host-microbe interaction genes, confirms that the plant immune system is a primary “gatekeeper” for microbial colonization (76–78). While individual ASV associations were sparse, treating microbial sub-communities as integrated units significantly increased the proportion of phenotypic variance explained. This suggests that complex agronomic traits are influenced by microbial consortia rather than isolated taxa. The co-selection of host traits and specific microbial networks suggests a joint host–microbiome basis for agronomic performance. Finally, the successful incorporation of microbiome data into genomic selection (GS) models—particularly for plant architecture and flowering—demonstrates a tangible path for microbiome-informed breeding. To transition these findings into routine practice, future research must prioritize functionally relevant and stable microbial predictors, validate these models across multi-environment trials, and develop high-throughput profiling methods.

As the first large-scale study of its kind in tobacco, this work outlines a reciprocal interplay between host genetics and the microbiome, establishing a framework for the next generation of microbiome-assisted crop improvement.

## Funding

This research was funded by the National Science Foundation of China (32200503), Agricultural Science and Technology Innovation Program (ASTIP-TRIC01) from Chinese Academy Agriculture Sciences, the Crop Germplasm Resources Protection and Utilization Special Grant from the Ministry of Agriculture and Rural Affairs of the People’s Republic of China, and the National Science and Technology Resource Sharing Service Platform (NCGRC-2023-18).

## Data Accessibility

## Conflict of Interest

The authors declare no conflict of interest.

## Notes

### Competing Interest Statement

The authors have declared no competing interest.

## Reference

1. Sul WJ. Host-Associated Microbiome. J Microbiol. 2024 Mar 1;62(3):135–6.

2. Chung IY, Kim J, Koh A. The Microbiome Matters: Its Impact on Cancer Development and Therapeutic Responses. J Microbiol. 2024 Mar 1;62(3):137–52.

3. Trivedi P, Leach JE, Tringe SG, Sa T, Singh BK. Plant–microbiome interactions: from community assembly to plant health. Nat Rev Microbiol. 2020 Nov 1;18(11):607–21.

4. Pieterse CMJ, Zamioudis C, Berendsen RL, Weller DM, Van Wees SCM, Bakker PAHM. Induced Systemic Resistance by Beneficial Microbes. Vol. 52, Annual Review of Phytopathology. Annual Reviews; 2014. p. 347–75.

5. Carrión VJ, Perez-Jaramillo J, Cordovez V, Tracanna V, de Hollander M, Ruiz-Buck D, et al. Pathogen-induced activation of disease-suppressive functions in the endophytic root microbiome. Science. 2019 Nov 1;366(6465):606–12.

6. Cheng YT, Zhang L, He SY. Plant-Microbe Interactions Facing Environmental Challenge. Cell Host Microbe. 2019 Aug 14;26(2):183–92.

7. Zahran Hamdi Hussein. Rhizobium-Legume Symbiosis and Nitrogen Fixation under Severe Conditions and in an Arid Climate. Microbiol Mol Biol Rev. 1999 Dec 1;63(4):968–89.

8. Backer R, Rokem JS, Ilangumaran G, Lamont J, Praslickova D, Ricci E, et al. Plant Growth-Promoting Rhizobacteria: Context, Mechanisms of Action, and Roadmap to Commercialization of Biostimulants for Sustainable Agriculture. Front Plant Sci [Internet]. 2018;Volume 9-2018. Available from: https://www.frontiersin.org/journals/plant-science/articles/10.3389/fpls.2018.01473

9. Singh BK, Trivedi P, Egidi E, Macdonald CA, Delgado-Baquerizo M. Crop microbiome and sustainable agriculture. Nat Rev Microbiol. 2020 Nov 1;18(11):601–2.

10. de Vries FT, Griffiths RI, Knight CG, Nicolitch O, Williams A. Harnessing rhizosphere microbiomes for drought-resilient crop production. Science. 2020 Apr 17;368(6488):270–4.

11. Agler MT, Ruhe J, Kroll S, Morhenn C, Kim ST, Weigel D, et al. Microbial Hub Taxa Link Host and Abiotic Factors to Plant Microbiome Variation. PLOS Biol. 2016 Jan 20;14(1):e1002352.

12. Brachi B, Filiault D, Whitehurst H, Darme P, Le Gars P, Le Mentec M, et al. Plant genetic effects on microbial hubs impact host fitness in repeated field trials. Proc Natl Acad Sci. 2022 July 26;119(30):e2201285119.

13. Peiffer JA, Spor A, Koren O, Jin Z, Tringe SG, Dangl JL, et al. Diversity and heritability of the maize rhizosphere microbiome under field conditions. Proc Natl Acad Sci. 2013 Apr 16;110(16):6548–53.

14. Edwards J, Johnson C, Santos-Medellín C, Lurie E, Podishetty NK, Bhatnagar S, et al. Structure, variation, and assembly of the root-associated microbiomes of rice. Proc Natl Acad Sci. 2015 Feb 24;112(8):E911–20.

15. Schlaeppi K, Dombrowski N, Oter RG, Ver Loren van Themaat E, Schulze-Lefert P. Quantitative divergence of the bacterial root microbiota in Arabidopsis thaliana relatives. Proc Natl Acad Sci. 2014 Jan 14;111(2):585–92.

16. Tian D, Wang P, Tang B, Teng X, Li C, Liu X, et al. GWAS Atlas: a curated resource of genome-wide variant-trait associations in plants and animals. Nucleic Acids Res. 2020 Jan 8;48(D1):D927–32.

17. Zhu X, Yang R, Liang Q, Yu Y, Wang T, Meng L, et al. Graph-based pangenome provides insights into structural variations and genetic basis of metabolic traits in potato. Mol Plant. 2025 Apr 7;18(4):590–602.

18. Wang W, Mauleon R, Hu Z, Chebotarov D, Tai S, Wu Z, et al. Genomic variation in 3,010 diverse accessions of Asian cultivated rice. Nature. 2018 May;557(7703):43–9.

19. Zhou Y, Zhang Z, Bao Z, Li H, Lyu Y, Zan Y, et al. Graph pangenome captures missing heritability and empowers tomato breeding. Nature. 2022 June 1;606(7914):527–34.

20. Yang L, He W, Zhu Y, Lv Y, Li Y, Zhang Q, et al. GWAS meta-analysis using a graph-based pan-genome enhanced gene mining efficiency for agronomic traits in rice. Nat Commun. 2025 Apr 3;16(1):3171.

21. Wang Y, Wang X, Sun S, Jin C, Su J, Wei J, et al. GWAS, MWAS and mGWAS provide insights into precision agriculture based on genotype-dependent microbial effects in foxtail millet. Nat Commun. 2022 Oct 7;13(1):5913.

22. Horton MW, Bodenhausen N, Beilsmith K, Meng D, Muegge BD, Subramanian S, et al. Genome-wide association study of Arabidopsis thaliana’s leaf microbial community. Nat Commun. 2014 Nov 10;5:5320.

23. He X, Wang D, Jiang Y, Li M, Delgado-Baquerizo M, McLaughlin C, et al. Heritable microbiome variation is correlated with source environment in locally adapted maize varieties. Nat Plants. 2024 Apr;10(4):598–617.

24. Deng S, Caddell DF, Xu G, Dahlen L, Washington L, Yang J, et al. Genome wide association study reveals plant loci controlling heritability of the rhizosphere microbiome. ISME J. 2021 Nov 1;15(11):3181–94.

25. Turpin W, Espin-Garcia O, Xu W, Silverberg MS, Kevans D, Smith MI, et al. Association of host genome with intestinal microbial composition in a large healthy cohort. Nat Genet. 2016 Nov 1;48(11):1413–7.

26. Neiverth A, Delai S, Garcia DM, Saatkamp K, de Souza EM, Pedrosa F de O, et al. Performance of different wheat genotypes inoculated with the plant growth promoting bacterium Herbaspirillum seropedicae. Eur J Soil Biol. 2014 Sept 1;64:1–5.

27. Rodriguez PA, Rothballer M, Chowdhury SP, Nussbaumer T, Gutjahr C, Falter-Braun P. Systems Biology of Plant-Microbiome Interactions. Mol Plant. 2019 June 3;12(6):804–21.

28. Peedin GF, Gerald F. Tobacco cultivation. Spec Crops. 2011;

29. Whitham S, McCormick S, Baker B. The N gene of tobacco confers resistance to tobacco mosaic virus in transgenic tomato. Proc Natl Acad Sci. 1996;93(16):8776–81.

30. An G. High Efficiency Transformation of Cultured Tobacco Cells 1. Plant Physiol. 1985 Oct 1;79(2):568–70.

31. Zhang J, Zhang Y, Du Y, Chen S, Tang H. Dynamic metabonomic responses of tobacco (Nicotiana tabacum) plants to salt stress. J Proteome Res. 2011;10(4):1904–14.

32. Yuan G, Sun K, Yu W, Jiang Z, Jiang C, Liu D, et al. Development of a MAGIC population and high-resolution quantitative trait mapping for nicotine content in tobacco. Front Plant Sci [Internet]. 2023;Volume 13-2022. Available from: https://www.frontiersin.org/journals/plant-science/articles/10.3389/fpls.2022.1086950

33. Shi R, Jin J, Nifong JM, Shew D, Lewis RS. Homoeologous chromosome exchange explains the creation of a QTL affecting soil-borne pathogen resistance in tobacco. Plant Biotechnol J. 2022 Jan 1;20(1):47–58.

34. Liu Y, Yuan G, Si H, Sun Y, Jiang Z, Liu D, et al. Identification of QTLs Associated With Agronomic Traits in Tobacco via a Biparental Population and an Eight-Way MAGIC Population. Front Plant Sci [Internet]. 2022;Volume 13-2022. Available from: https://www.frontiersin.org/journals/plant-science/articles/10.3389/fpls.2022.878267

35. Zan Y, Chen S, Ren M, Liu G, Liu Y, Han Y, et al. The genome and GeneBank genomics of allotetraploid Nicotiana tabacum provide insights into genome evolution and complex trait regulation. Nat Genet. 2025 Apr 1;57(4):986–96.

36. Callahan BJ, McMurdie PJ, Rosen MJ, Han AW, Johnson AJA, Holmes SP. DADA2: High-resolution sample inference from Illumina amplicon data. Nat Methods. 2016 July 1;13(7):581–3.

37. Bolyen E, Rideout JR, Dillon MR, Bokulich NA, Abnet CC, Al-Ghalith GA, et al. Reproducible, interactive, scalable and extensible microbiome data science using QIIME 2. Nat Biotechnol. 2019 Aug 1;37(8):852–7.

38. Bokulich NA, Kaehler BD, Rideout JR, Dillon M, Bolyen E, Knight R, et al. Optimizing taxonomic classification of marker-gene amplicon sequences with QIIME 2’s q2-feature-classifier plugin. Microbiome. 2018 May 17;6(1):90.

39. Yilmaz P, Parfrey LW, Yarza P, Gerken J, Pruesse E, Quast C, et al. The SILVA and “all-species living tree project (LTP)” taxonomic frameworks. Nucleic Acids Res. 2014;42(D1):D643–8.

40. Oksanen J. Vegan: community ecology package version 1.8-6. Httpcran R-Proj Org. 2007;

41. Segata N, Izard J, Waldron L, Gevers D, Miropolsky L, Garrett WS, et al. Metagenomic biomarker discovery and explanation. Genome Biol. 2011 June 24;12(6):R60.

42. Li H. Aligning sequence reads, clone sequences and assembly contigs with BWA-MEM. ArXiv Prepr ArXiv13033997. 2013;

43. Danecek P, Bonfield JK, Liddle J, Marshall J, Ohan V, Pollard MO, et al. Twelve years of SAMtools and BCFtools. GigaScience. 2021 Feb 1;10(2):giab008.

44. Garrison E, Marth G. Haplotype-based variant detection from short-read sequencing. ArXiv Prepr ArXiv12073907. 2012;

45. Danecek P, Auton A, Abecasis G, Albers CA, Banks E, DePristo MA, et al. The variant call format and VCFtools. Bioinformatics. 2011 Aug 1;27(15):2156–8.

46. Ayres DL, Darling A, Zwickl DJ, Beerli P, Holder MT, Lewis PO, et al. BEAGLE: An Application Programming Interface and High-Performance Computing Library for Statistical Phylogenetics. Syst Biol. 2012 Jan 1;61(1):170–3.

47. Yang J, Lee SH, Goddard ME, Visscher PM. GCTA: A Tool for Genome-wide Complex Trait Analysis. Am J Hum Genet. 2011 Jan 7;88(1):76–82.

48. Liu C, Cui Y, Li X, Yao M. microeco: an R package for data mining in microbial community ecology. FEMS Microbiol Ecol. 2021;97(2):fiaa255.

49. Wallace C. A more accurate method for colocalisation analysis allowing for multiple causal variants. PLoS Genet. 2021;17(9):e1009440.

50. Zhang F, Chen W, Zhu Z, Zhang Q, Nabais MF, Qi T, et al. OSCA: a tool for omic-data-based complex trait analysis. Genome Biol. 2019 May 28;20(1):107.

51. Palberg D, Kisiała A, Jorge GL, Emery RJN. A survey of Methylobacterium species and strains reveals widespread production and varying profiles of cytokinin phytohormones. BMC Microbiol. 2022 Feb 8;22(1):49.

52. Krug L, Morauf C, Donat C, Müller H, Cernava T, Berg G. Plant Growth-Promoting Methylobacteria Selectively Increase the Biomass of Biotechnologically Relevant Microalgae. Front Microbiol [Internet]. 2020;Volume 11-2020. Available from: https://www.frontiersin.org/journals/microbiology/articles/10.3389/fmicb.2020.00427

53. Sun H, Chen M, Wei L, Xue P, Zhao Q, Gao P, et al. Roots recruited distinct rhizo-microbial communities to adapt to long-term Cd and As co-contaminated soil in wheat-maize rotation. Environ Pollut. 2024 Feb 1;342:123053.

54. Zou Z, Yuan K, Ming L, Li Z, Yang Y, Yang R, et al. Changes in Alpine Soil Bacterial Communities With Altitude and Slopes at Mount Shergyla, Tibetan Plateau: Diversity, Structure, and Influencing Factors. Front Microbiol [Internet]. 2022;Volume 13-2022. Available from: https://www.frontiersin.org/journals/microbiology/articles/10.3389/fmicb.2022.839499

55. Wang Y, Du L, Liu H, Long D, Huang M, Wang Y, et al. Halosulfuron methyl did not have a significant effect on diversity and community of sugarcane rhizosphere microflora. J Hazard Mater. 2020 Nov 15;399:123040.

56. Chen L, Hao Z, Li K, Sha Y, Wang E, Sui X, et al. Effects of growth-promoting rhizobacteria on maize growth and rhizosphere microbial community under conservation tillage in Northeast China. Microb Biotechnol. 2021 Mar 1;14(2):535–50.

57. Chen L, Li K, Shang J, Wu Y, Chen T, Wanyan Y, et al. Plant growth–promoting bacteria improve maize growth through reshaping the rhizobacterial community in low-nitrogen and low-phosphorus soil. Biol Fertil Soils. 2021 Nov 1;57(8):1075–88.

58. Jin M, Liu H, Liu X, Guo T, Guo J, Yin Y, et al. Complex genetic architecture underlying the plasticity of maize agronomic traits. Plant Commun. 2023 May 8;4(3):100473.

59. Compant S, Samad A, Faist H, Sessitsch A. A review on the plant microbiome: Ecology, functions, and emerging trends in microbial application. Spec Issue Plant Microbiome. 2019 Sept 1;19:29–37.

60. Xiong C, Zhu YG, Wang JT, Singh B, Han LL, Shen JP, et al. Host selection shapes crop microbiome assembly and network complexity. New Phytol. 2021 Jan 1;229(2):1091–104.

61. Chen P, Zhao M, Tang F, Hu Y, Peng X, Shen S. The effect of plant compartments on the Broussonetia papyrifera-associated fungal and bacterial communities. Appl Microbiol Biotechnol. 2020 Apr 1;104(8):3627–41.

62. Cregger MA, Veach AM, Yang ZK, Crouch MJ, Vilgalys R, Tuskan GA, et al. The Populus holobiont: dissecting the effects of plant niches and genotype on the microbiome. Microbiome. 2018 Feb 12;6(1):31.

63. Zarraonaindia Iratxe, Owens Sarah M., Weisenhorn Pamela, West Kristin, Hampton-Marcell Jarrad, Lax Simon, et al. The Soil Microbiome Influences Grapevine-Associated Microbiota. mBio. 2015 Mar 24;6(2):10.1128/mbio.02527-14.

64. Zhu YG, Peng J, Chen C, Xiong C, Li S, Ge A, et al. Harnessing biological nitrogen fixation in plant leaves. Trends Plant Sci. 2023 Dec 1;28(12):1391–405.

65. Chen S, Waghmode TR, Sun R, Kuramae EE, Hu C, Liu B. Root-associated microbiomes of wheat under the combined effect of plant development and nitrogen fertilization. Microbiome. 2019 Oct 22;7(1):136.

66. Flores-Félix JD, Menéndez E, Rivera LP, Marcos-García M, Martínez-Hidalgo P, Mateos PF, et al. Use of Rhizobium leguminosarum as a potential biofertilizer for Lactuca sativa and Daucus carota crops. J Plant Nutr Soil Sci. 2013 Dec 1;176(6):876–82.

67. Asaf S, Numan M, Khan AL, Al-Harrasi A. Sphingomonas: from diversity and genomics to functional role in environmental remediation and plant growth. Crit Rev Biotechnol. 2020 Feb 17;40(2):138–52.

68. Kim YJ, Park JY, Balusamy SR, Huo Y, Nong LK, Thi Le H, et al. Comprehensive Genome Analysis on the Novel Species Sphingomonas panacis DCY99T Reveals Insights into Iron Tolerance of Ginseng. Int J Mol Sci. 2020;21(6):2019.

69. Severino R, Froufe HJC, Barroso C, Albuquerque L, Lobo-da-Cunha A, da Costa MS, et al. High-quality draft genome sequence of Gaiella occulta isolated from a 150 meter deep mineral water borehole and comparison with the genome sequences of other deep-branching lineages of the phylum Actinobacteria. MicrobiologyOpen. 2019 Sept 1;8(9):e00840.

70. Kalam S, Basu A, Ahmad I, Sayyed RZ, El-Enshasy HA, Dailin DJ, et al. Recent Understanding of Soil Acidobacteria and Their Ecological Significance: A Critical Review. Front Microbiol [Internet]. 2020;Volume 11-2020. Available from: https://www.frontiersin.org/journals/microbiology/articles/10.3389/fmicb.2020.580024

71. Wei G, Ning K, Zhang G, Yu H, Yang S, Dai F, et al. Compartment Niche Shapes the Assembly and Network of Cannabis sativa-Associated Microbiome. Front Microbiol [Internet]. 2021;Volume 12-2021. Available from: https://www.frontiersin.org/journals/microbiology/articles/10.3389/fmicb.2021.714993

72. Philippot L, Raaijmakers JM, Lemanceau P, van der Putten WH. Going back to the roots: the microbial ecology of the rhizosphere. Nat Rev Microbiol. 2013 Nov 1;11(11):789–99.

73. Tipayno SC, Truu J, Samaddar S, Truu M, Preem JK, Oopkaup K, et al. The bacterial community structure and functional profile in the heavy metal contaminated paddy soils, surrounding a nonferrous smelter in South Korea. Ecol Evol. 2018 June 1;8(12):6157–68.

74. Compant Stéphane, Duffy Brion, Nowak Jerzy, Clément Christophe, Barka Essaïd Ait. Use of Plant Growth-Promoting Bacteria for Biocontrol of Plant Diseases: Principles, Mechanisms of Action, and Future Prospects. Appl Environ Microbiol. 2005 Sept 1;71(9):4951–9.

75. Roman-Reyna V, Pinili D, Borja FN, Quibod IL, Groen SC, Mulyaningsih ES, et al. The rice leaf microbiome has a conserved community structure controlled by complex host-microbe interactions. bioRxiv. 2019 Jan 1;615278.

76. Liu Q, Cheng L, Nian H, Jin J, Lian T. Linking plant functional genes to rhizosphere microbes: a review. Plant Biotechnol J. 2023 May 1;21(5):902–17.

77. Fitzpatrick CR, Salas-González I, Conway JM, Finkel OM, Gilbert S, Russ D, et al. The Plant Microbiome: From Ecology to Reductionism and Beyond. Vol. 74, Annual Review of Microbiology. Annual Reviews; 2020. p. 81–100.

78. Wippel K, Tao K, Niu Y, Zgadzaj R, Kiel N, Guan R, et al. Host preference and invasiveness of commensal bacteria in the Lotus and Arabidopsis root microbiota. Nat Microbiol. 2021 Sept 1;6(9):1150–62.

